# Covalent Targeting of Glutamate Cysteine Ligase to Inhibit Glutathione Synthesis

**DOI:** 10.1101/2023.05.18.541312

**Authors:** Lydia Zhang, Michelle Tang, Xavier Tao, Qian Shao, Vienna Thomas, Saki Shimizu, Miki Kasano, Yoshinori Ishikawa, Takayuki Inukai, Daniel K. Nomura

## Abstract

Dysregulated oxidative stress plays a major role in cancer pathogenesis and some types of cancer cells are particularly vulnerable to inhibiting cellular antioxidant capacity. Glutamate-cysteine ligase (GCL) is the first and rate-limiting step in the synthesis of the major cellular antioxidant glutathione (GSH). Developing a GCL inhibitor may be an attractive therapeutic strategy for certain cancer types that are particularly sensitive to oxidative stress. In this study, we reveal a cysteine-reactive covalent ligand EN25 that covalently targets an allosteric cysteine C114 on GCLM, the modifier subunit of GCL, leading to inhibition of GCL activity, lowering of cellular GSH levels, and impaired cell viability in ARID1A-deficient cancer cells that are particularly vulnerable to glutathione depletion, but not in ARID1A-positive cancer cells. Our studies uncover a novel potential ligandable site within GCLM that can be targeted to inhibit the GSH synthesis in cancer cells to target vulnerable cancer cell types.

---

Many different types of cancers have dysregulated redox homeostasis which contributes to fueling cancer pathogenicity, but also presents a vulnerability that can be pharmacologically targeted for cancer therapy. For example, MYC-driven liver tumors show suppression of glutathione synthesis, and instead use glutamine-derived carbons for used for other metabolic pathways to support tumor growth. However, this alteration in metabolism makes these MYC-driven tumors hyper-sensitive to oxidative stress ^[1]^. Previous studies have revealed that ARID1A-deficient cancers, through their downregulation of the cystine transporter SLC7A11, are specifically vulnerable to inhibition of glutathione synthesis ^[2]^. Thus, targeting the synthesis of glutathione in cancer cells is an attractive strategy towards impairing certain types of human cancers through heightening oxidative stress and potentially inducing ferroptosis-mediated cell death in cancer cells ^[3]^.

Glutamate-cysteine ligase (GCL) consists of two subunits--glutamate-cysteine ligase modifier subunit (GCLM) and glutamate-cysteine ligase catalytic subunit (GCLC)—and are two enzymes involved in the synthesis of glutathione (GSH) ^[4]^. While developing GCL inhibitors would potentially have chemotherapeutic implications, there are no drug-like GCL inhibitors. Buthionine sulfoximine (BSO), a substrate mimetic with poor drug-like properties, currently represents one of the best currently available GCL inhibitors ^[5]^. In this study, we sought to use covalent ligand discovery approaches to identify unique ligandable sites within and functional inhibitors for GCL.

We performed a screen with a library of 679 cysteine-reactive covalent ligands in a GCL activity screen to identify covalent inhibitors of GCL activity **(Fig. 1a, Table S1)**. Through this screen, we identified the acrylamide EN25 that dose-responsively inhibited GCL activity with a 50% inhibitory concentration (IC_50_) of 16 μM **(Fig. 1a-1c)**. We next sought to identify the site of EN25 modification on the GCL protein complex by performing LC-MS/MS analysis of tryptic digests from GCLC and GCLM labeled with EN25. We identified selective and covalent EN25 modification of C114 on GCLM **(Fig. 1d)**. Given that GCLM is the regulatory subunit of the GCL complex, we postulate that covalent targeting of C114 results in allosteric inhibition of GCLC catalytic enzyme activity. To further demonstrate covalent modification of this scaffold with GCLM, we synthesized an alkyne-functionalized analog of EN25 with EN25-alkyne that demonstrated an IC_50_ for GCL activity of 3.6 μM. This probe showed covalent labeling of GCLM and dose-responsive competition of this labeling by EN25 **(Fig. 1e-1f)**. To understand the proteome-wide selectivity of EN25, we next performed cysteine chemoproteomic profiling using isotopic tandem orthogonal proteolysis-enabled activity-based protein profiling (isoTOP-ABPP) ^[6,7]^. We treated A2780 ARID1A-negative cancer cells with either vehicle or EN25, and subsequently treated resulting cell lysates with the alkyne functionalized cysteine-reactive iodoacetamide probe, after which isotopically encoded cleavable biotin-azide enrichment handles were appended onto probe-labeled proteins by click-chemistry, and subjected to avidin-enrichment, tryptic digestion, elution of probe-modified tryptic peptides, and analysis by quantitative proteomic methods. While we were not able to identify a probe-modified tryptic peptide encompassing C114 of GCLM, we identified nine potential off-targets of EN25 out of >5000 cysteines within proteins quantified in cells **(Fig. S1; Table S2)**. These nine targets included HNRNPA3 C85, MTHFD1L C779, C9Orf142 C180, SIRT6 C18, GPKOW C137, PNN C439, ALDH7A1 C478, NUP85 C515, and U2URP C62. None of these nine off-targets were likely to influence glutathione metabolism in cancer cells, and we deemed this overall selectivity to be sufficient for follow-up studies.

**Figure 1.**
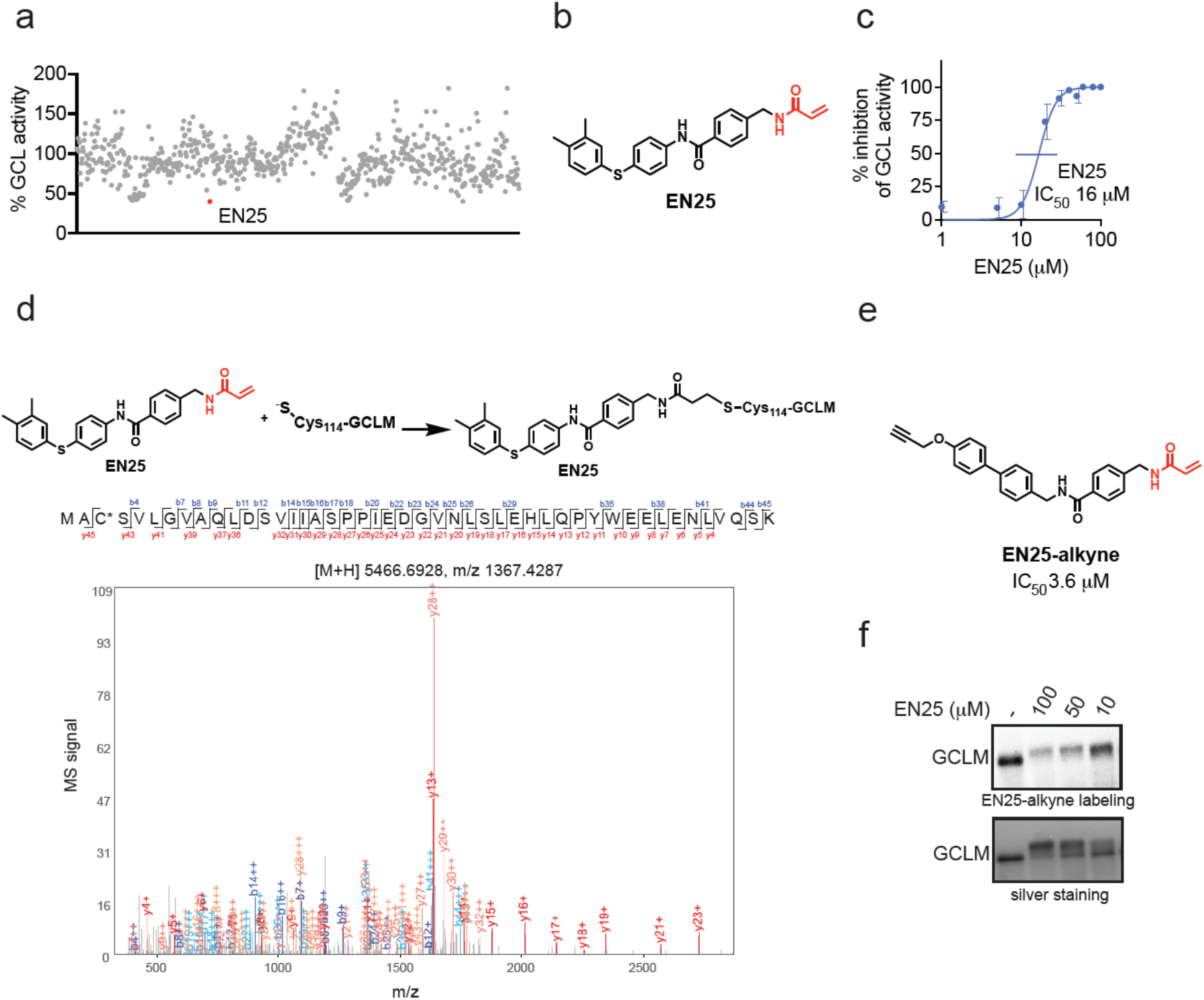
Identification of a covalent inhibitor of the GCL complex. **(a)** A functional screen of cysteine-reactive covalent ligands against GCL to look for inhibition of GCL activity. Pure GCL protein was incubated with its substrates, cofactors, and DMSO vehicle or covalent ligand. The product, gamma-glutamylcysteine (GluCys), was converted to a derivative of 2,3-napthalenedicarboxaldehyde (NDA), NDA-GluCys, to enable a colorimetric readout of product formation. The top hit, EN25, is shown in red. **(b)** Structure of EN25 with the cysteine-reactive acrylamide moiety highlighted in red. **(c)** EN25 showed an IC_50_ of 16 μM on GCL activity. **(d)** MS/MS analysis of EN25 adduct on the GCL complex showed that EN25 modifies C114 on GCLM. Pure human GCLM and GCLC were incubated with EN25 (50μM) for 30 min at 37°C. The GCL complex was then subjected to tryptic digestion, and tryptic digests were analyzed by LC-MS/MS. **(e)** Structure of the alkyne-functionalized analog of EN25 (EN25-alkyne) that demonstrated an IC_50_ of 3.6 μM on GCL activity. **(f)** Gel-based ABPP showing that EN25 displaces EN25-alkyne labeling of pure human GCLM protein. GCLM was pre-incubated with DMSO vehicle or EN25 for 30 min at 37°C, prior to EN25-alkyne labeling (50 μM final) for 2 h at 37°C. Rhodamine-azide was appended to probe-modified protein by CuAAC, subjected to SDS/PAGE, and analyzed by in-gel fluorescence. Shown below is a silver stain of the gel to show equal protein loading. Graph shown in **(a)** is representative of n=2 biologically independent replicates/group and percentage values for each compound and structures of compounds screened are listed in **Table S1**. Graph shown in **(c)** is representative of n=4 biologically independent replicates/group. Gel shown in **(f)** is representative of n=3 biologically independent replicates/group.

Given that ARID1A-negative cancer cells have been previously shown to be more susceptible to inhibition of glutathione synthesis compared to ARID1A-positive cancer cells, we next sought to determine whether EN25 selectively impairs cell viability of ARID1A-negative A2780 ovarian cancer cells, compared to ARID1A-positive RMG-1 ovarian cancer cells. Consistent with this premise, RMG-1 cancer cells showed resistance to EN25-mediated cell viability impairments compared to the more sensitive A2780 cancer cell line **(Fig. 2a)**. We further demonstrated that EN25 significantly lowered GSH levels in A2780 cancer cells compared to no changes observed in RMG-1 cells **(Fig. 2b)**. To confirm that the effects we observed were through on-target activity, we further demonstrated that GCLM knockdown with siRNA conferred hypersensitivity to EN25-mediated cell viability impairments compared to non-targeting siControl A2780 cells **(Fig. 2c-2d)**.

**Figure 2.**
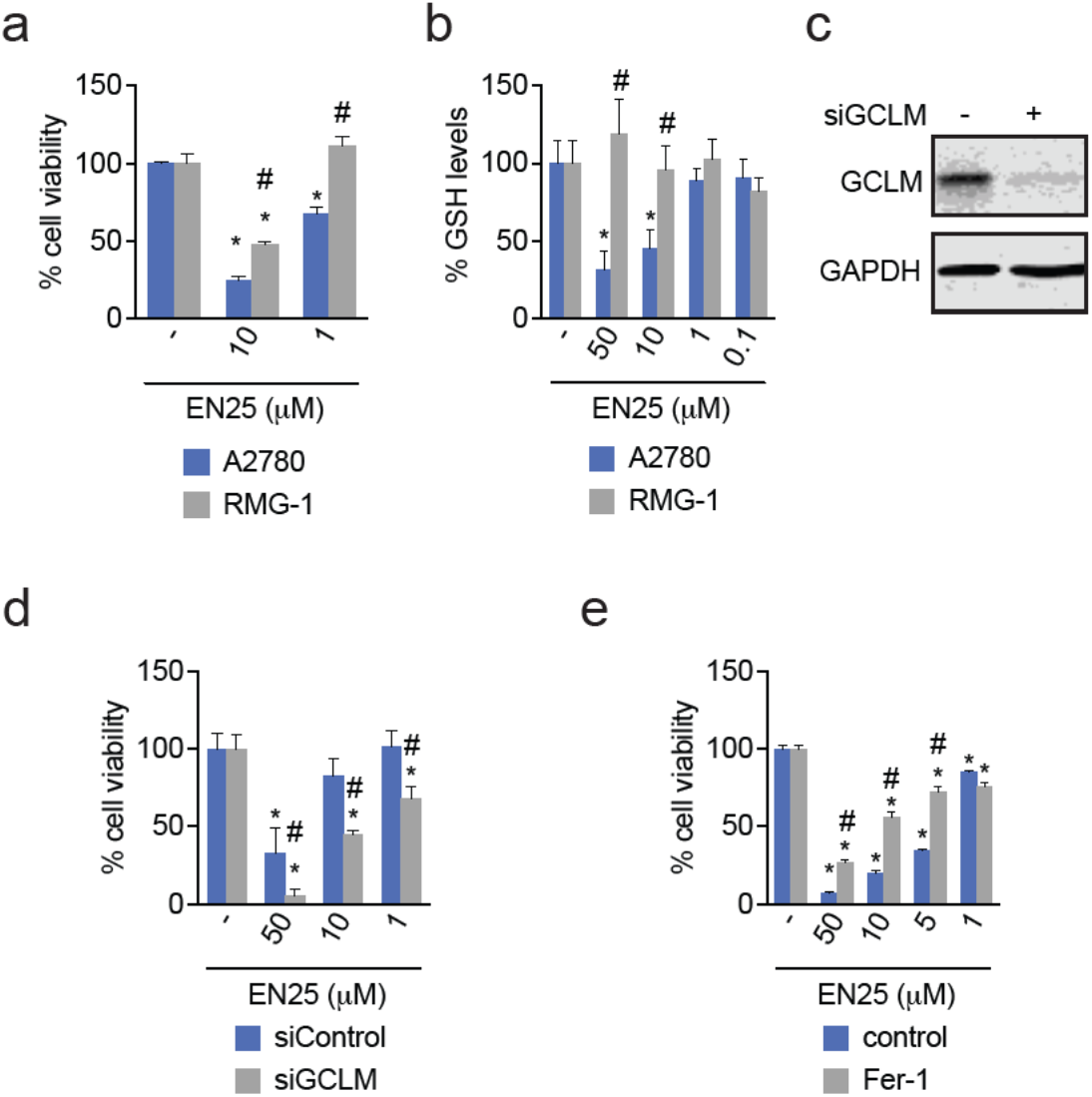
Characterizing the effect of EN25 on GCLM and glutathione (GSH) levels in vitro. **(a)** EN25 impairs A2780 cell viability over RMG-1 cell viability. Cells were treated with DMSO vehicle or EN25 and cell proliferation was assessed by Hoechst staining at 24 hr. **(b)** EN25 selectively reduces GSH levels in A2780 over RMG-1 cell lines. Cells were treated with DMSO vehicle or EN25 and GSH levels were detected by single reaction monitoring (SRM) LC-MS/MS 4hr post treatment. **(c-d)** GCLM knockdown with siRNA in A2780 cells confers hypersensitivity to EN25-mediated cell viability impairments. A2780 cells were transfected with siControl or siGCLM for transient knockdown and used for experiments 48 hr after transfection. **(c)** Western blot for GCLM and loading control GAPDH in transfected A2780 cells. **(d)** Cell proliferation of siGCLM A2780 cell lines compared to siControl A2780 cell lines after 24 hr treatment with EN25. Proliferation was measured by Hoechst staining. **(e)** Ferrostatin-1 attenuates EN25-mediated cell viability impairments in A2780. A2780 cells were treated with DMSO vehicle or 2 μM of Ferrostatin-1 immediately prior to DMSO vehicle or EN25 treatment. Cell viability was measured at 48 hr with Hoechst staining. Data in **(a, b, c, d, e)** are shown as percentage compared with DMSO vehicle control and show average ± sem of n=6 for **(a, c, d, e)** and n=3 for **(b)** biologically independent replicates/group. Statistical significance is calculated as *p < 0.05 compared to DMSO vehicle and #p < 0.05 compared to cells treated with EN25 alone. Blot shown in **(c)** is representative of n=3 biologically independent replicates/group.

We next hypothesized that GCL inhibition and depletion of cellular GSH levels with EN25 treatment in ARID1A-negative cancer cells would lead to accumulation of oxidative stress, lipid peroxidation, and ferroptosis-mediated cell death. Consistent with this premise, a ferroptosis inhibitor ferrostatin attenuated the cell viability impairments conferred by EN25 treatment in A2780 cancer cells **(Fig. 2e)**. These results indicate that EN25 likely impairs cancer cell proliferation through ferroptosis-mediated mechanisms. We also explored structure-activity relationship of EN25 analogs that were commercially available. Through testing of 12 EN25 analogs that were commercially obtained through Enamine, we found that EN25-7 and EN25-8 showed the best inhibitory potency against GCL with IC50s of 8.9 and 6.8 μM, respectively **(Fig. 3)**.

**Figure 3.**
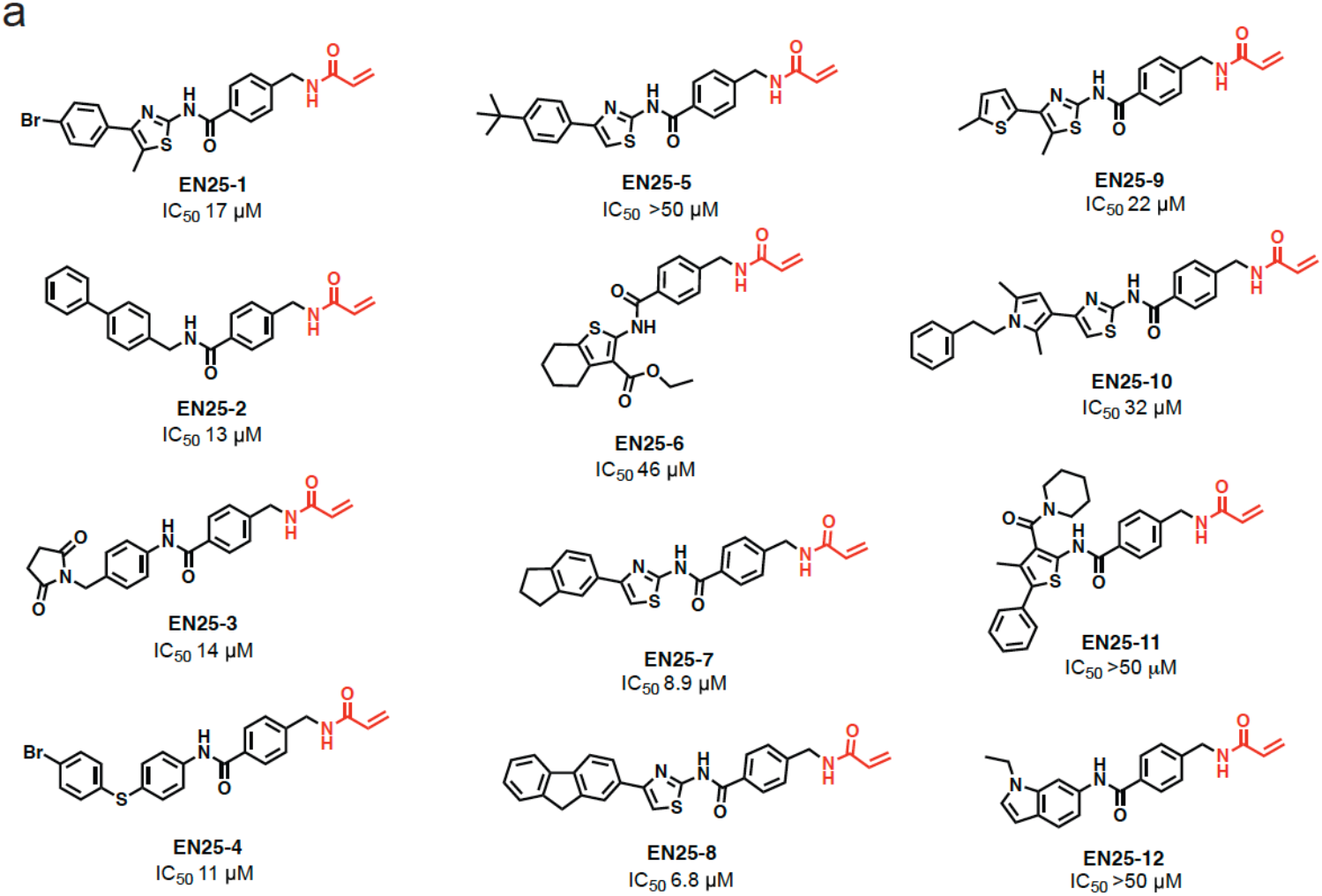
Structures of EN25 analogs and associated IC_50_ values for GCL inhibition. **(a)** Structures of EN25 analogs with the cysteine reactive acrylamide moieties highlighted in red. The associated IC_50_ values listed below were determined via the functional assay described in **(Fig. 1a)**.

Overall, in this study, we have discovered novel GCL inhibitors that uniquely inhibit GCL activity through allosteric and covalent targeting of C114 on the regulatory GCLM subunit. We demonstrate that the GCL inhibitor EN25 inhibits GCL activity, lowers glutathione levels and impairs cancer cell proliferation, preferentially in ARID1A-negative cancer cells through a ferroptosis-mediated mechanism. C114 on GCLM and EN25 represent novel ligands and ligandable sites for inhibiting GCL activity.

## Methods

### Materials

Cysteine-reactive covalent ligand libraries purchased from Enamine or their synthesis were previously described ^[8–11]^. Their structures are all shown in **Table S1**.

### Cell Culture

A2780 cells (obtained from Sigma-Aldrich, 93112519-1VL) were cultured in RPMI-1640 containing 10% (v/v) fetal bovine serum (FBS) and 10mM glutamine and maintained at 37°C with 5% CO2. RMG-1 cells (obtained from Fisher Scientific, NC0992912) were cultured in Ham’s F12 Medium containing 10% (v/v) fetal bovine serum (FBS) and 10mM glutamine and maintained at 37°C with 5% CO2.

### GCL Functional Assay Screen

GCL activity was determined by the fluorescence assay described by Chen *et al*.^[12]^ For the activity assay screen, pure human recombinant GCLC (0.05μM) and GCLM (0.1μM) protein (obtained from Ono Pharmaceuticals) were mixed in PBS and aliquoted into a 96-well v-bottom assay plate (Corning, P-96-450V-C) in a total volume of 20μl. Covalent ligands or DMSO vehicle were then added (0.5 μl) to the protein and incubated for 30 minutes at 37ºC. Next, 20 μl of GCL-reaction cocktail [400 mM Tris-HCl, 40 mM adenosine triphosphate (ATP), 40 mM L-glutamic acid, 2 mM EDTA, 20 mM sodium borate, 2 mM serine and 40 mM MgCl_2_] was added to each well and incubated at 37°C for 5 min. The reaction was initiated with 20 μl of 30 mM cysteine solution (dissolved in TES/SB buffer) and the mixtures were incubated at 37°C for 15 min. The GCL reaction was stopped by precipitation of proteins with 200 mM 5-sulfosalicylic acid (Sigma Aldrich, 247006) and then put onto ice for 20 min. Afterwards, the mixtures were centrifuged at 2,500 rpm at 4°C for 10 min and 20 μl of the supernatant that contained the γ-GC product was added to a half-area black 96-well plate (Corning, 3904). For fluorescence detection, 100 μl 2,3-naphthalenedicarboxyaldehyde (NDA) solution [50 mM Tris, pH 10, 0.5 N NaOH, and 10 mM NDA in Me2SO, v/v/v 1.4/0.2/0.2] was added into each well and incubated in the dark at room temperature for 30 min. The formation of NDA-γ-GC was measured (472 nm excitation/528 nm emission) using a fluorescent plate reader (Spark; Tecan Group Ltd.)

### LC/MS-MS Analysis of EN25 Interactions with GCLC/GCLM

The GCL complex made in a 2:1 GCLM:GCLC molar ratio totaling 50 μg in 50 μL PBS was incubated for 30 min at 37°C with EN25 (50 μM). The sample was precipitated by addition of 10 μL of 100% (w/v) TCA and cooled to -80 °C for 1 h. The sample was then spun at max speed for 10 min at 4°C and washed three times with ice cold 0.01 M HCl/90 % acetone solution. The pellet was resuspended in 4 M urea containing 0.1 % Protease Max (Promega Corp. V2071) and diluted in 40 mM ammonium bicarbonate buffer. Next, the sample was reduced with 10μl of 10 mM TCEP and incubated at 60 °C for 30 min. Afterwards, 10μl of 150mM iodoacetamide was added and incubated at room temperature for 30 minutes. The sample was then diluted 50% with PBS before addition of sequencing grade trypsin (1 μg per sample, Promega Corp, V5111) and incubated overnight at 37 °C. Samples were acidified to a final concentration of 5 % formic acid and centrifuged at 13,200 rpm for 30 min. The supernatant was transferred to a new tube and stored at -80°C until MS analysis and were analyzed as previously described, searching for the predicted mass adduct on cysteines ^[11,13]^.

### Cell Viability Assessment

Cell viability assays were performed using Hoechst 33342 dye (Invitrogen, H3570) according to the manufacturer’s protocol and as previously described ^[10]^. Cells were seeded into 96-well plates (6,000 per well) in a total volume of 200 μL and left to adhere overnight. Cells were treated with 1 μL of compound stock or DMSO vehicle. After the appropriate incubation period, media was aspirated from each well and 100 μL of staining solution containing 10% formalin and Hoechst 33342 dye was added to each well and incubated in the dark for 15 min at room temperature. Staining solution was removed, and wells were washed three times with PBS before fluorescent reading.

### LC-MS/MS-Based Quantification of GSH Levels in Cells

Cells were seeded into 6 cm dishes at 2x10^6^ million cells per well in 2 mL and allowed to adhere overnight. Cells were treated the next day with DMSO or EN25 for 4 h before harvesting on ice with PBS. Cells were lysed by probe sonication and pelleted by centrifugation. Pellets were frozen at -80°C for one hour. For polar extraction, 180μl of 40:40:20 acetonitrile:methanol:water was added to each frozen cell pellet on dry ice. An internal standard of L-Serine-^15^N,d_3_ (1 nmole final, Cambridge Isotope Labs, DNLM-6863) was added to each sample. Precipitated proteins and lipids were pelleted by centrifugation at 16,000 x g for 10 min. The supernatant was then transferred into a polyspring insert inside an LC-MS/MS vial, briefly vortexed, and ran through a QQQ mass spec through a C18 reverse phase column (Phenomenex, 00B-4435-E0) at a 0.7ml/min flow rate. GSH levels were detected by single reaction monitoring (SRM) and normalized to internal standard and compared against GSH levels in DMSO-treated controls. Data was analyzed as previously reported ^[14]^.

### Determining GSH Levels using the GSH-Glo Kit

To assess cellular glutathione levels, we used the GSH-Glo™ Glutathione Assay Kit (Promega, V6911) and the protocol was followed as described in the manufacture’s instructions. Cells were seeded into flat white 96-well plates (Corning, 3917) at 6,000 cells per well in 200 μL and allowed to adhere overnight. Cells were then treated with DMSO vehicle or compound. After the appropriate treatment time, media was aspirated from each well before addition of 1X GSH-Glo™ Reagent (100 μl) and incubated at room temperature for 30 minutes on a plate shaker. Next, Luciferin Detection Reagent (100 μl) was added to the wells and the plate was incubated at room temperature for 15 minutes on a plate shaker before measurement of luminescence with a plate reader.

### Gel-based ABPP

Gel-based ABPP methods were performed as previously described ^[11]^. Recombinant pure human GCLM (0.1 μg), provided by from Ono Pharmaceuticals, was pre-treated with DMSO or the covalently-acting EN25-alkyne probe (10-100μM final concentration) for 30 minutes at 37 °C in a volume of 50μL of PBS. CuAAC was performed to append rhodamine-azide (1 μM final concentration) onto alkyne probe-labeled proteins. Samples were then diluted with 30 μL of 4 x reducing Laemmli SDS sample loading buffer (Alfa Aesar) and heated at 90 °C for 5 min. The samples were separated on precast 4-20% TGX gels (Bio-Rad Laboratories, Inc.).

### IsoTOP-ABPP Chemoproteomic Profiling

IsoTOP-ABPP studies were done as previously reported ^[6,7,10,15]^. Cells were seeded and grown to a confluency of approximately 90% and treated for 90 min with either DMSO vehicle or EN25 before harvest and lysis. Cells were lysed by probe sonication in PBS and protein concentrations were measured by a bicinchoninic acid (BCA) assay and normalized to 4 mg/ml. Proteomes were subsequently labeled with 100 μM of a cysteine-reactive alkyne functionalized iodoacetamide probe (IA-alkyne) for 1 hour at room temperature. Copper-catalyzed azide-alkyne cycloaddition (CuAAC) was performed by sequential addition of 1 mM TCEP (Sigma), 34mM tris((1-benzyl-1H-1,2,3-triazol-4-yl)methyl)amine, 1mM copper(II) sulfate, and a TEV-cleavable biotin-linker-azide functionalized with an isotopically light (for control) or heavy (for treatment) valine. Following CuAAC, proteomes were precipitated by centrifugation at 6500 x g, washed in ice-cold methanol, combined in a 1:1 control/treated ratio, washed again, then denatured and resolubilized by heating in 1.2 % SDS/PBS to 80°C for 5 minutes. Insoluble components were precipitated by centrifugation at 6500 x g and the soluble proteome was diluted in 5 ml 0.2% SDS/PBS. Labeled proteins were bound to 170μl of avidin-agarose beads (Thermo Pierce, PI20349) per sample while rotating overnight at 4°C. Bead-linked proteins were enriched by washing three times each in PBS and water, then resuspended in 6 M urea/PBS (Sigma) and reduced in 1 mM TCEP, alkylated with 18 mM iodoacetamide (IA) (Sigma, 786-228), then washed and resuspended in 2 M urea and trypsinized overnight with 0.5 μg/μl sequencing grade trypsin (Promega, V5111). Tryptic peptides were eluted off with spin columns (BioRad, 7326204) under a vacuum manifold. Beads were then washed three times each in PBS and water, washed in TEV buffer solution (water, TEV buffer, 100 μM dithiothreitol), and resuspended in buffer with Ac-TEV protease and incubated overnight. Peptides were diluted in water and acidified with 1.2M formic acid (Thomas Scientific, A11750) in preparation for mass spectrometry analysis.

### IsoTOP-ABPP Mass Spectrometry Analysis

Peptides from all chemoproteomic experiments were pressure-loaded onto a 250 μm inner diameter fused silica capillary tubing packed with 4 cm of Aqua C18 reverse-phase resin (Phenomenex, 04A-4299), which was previously equilibrated on an Agilent 600 series high-performance liquid chromatograph using the gradient from 100% buffer A to 100% buffer B over 10 min, followed by a 5 min wash with 100% buffer B and a 5 min wash with 100% buffer A. The samples were then attached using a MicroTee PEEK 360 μm fitting (Thermo Fisher Scientific p-888) to a 13 cm laser pulled column packed with 10 cm Aqua C18 reverse-phase resin and 3 cm of strong-cation exchange resin for isoTOP-ABPP studies. Samples were analyzed using an Q Exactive Plus mass spectrometer (Thermo Fisher Scientific) using a five-step Multidimensional Protein Identification Technology (MudPIT) program, using 0, 25, 50, 80 and 100% salt bumps of 500 mM aqueous ammonium acetate and using a gradient of 5–55% buffer B in buffer A (buffer A: 95:5 water:acetonitrile, 0.1% formic acid; buffer B 80:20 acetonitrile:water, 0.1% formic acid). Data were collected in data-dependent acquisition mode with dynamic exclusion enabled (60 s). One full mass spectrometry (MS1) scan (400–1,800 mass-to-charge ratio (m/z)) was followed by 15 MS2 scans of the nth most abundant ions. Heated capillary temperature was set to 200 °C and the nanospray voltage was set to 2.75 kV. Data were extracted in the form of MS1 and MS2 files using Raw Extractor v.1.9.9.2 (Scripps Research Institute) and searched against the Uniprot human database using ProLuCID search methodology in IP2 v.3v.5 (Integrated Proteomics Applications, Inc.) (Xu et al., 2015). Cysteine residues were searched with a static modification for carboxyaminomethylation (+57.02146) and up to two differential modifications for methionine oxidation and either the light or heavy TEV tags (+464.28596 or +470.29977, respectively). Peptides were required to be fully tryptic peptides and to contain the TEV modification. ProLUCID data were filtered through DTASelect to achieve a peptide false-positive rate below 5%. Only those probe-modified peptides that were evident across two out of three biological replicates were interpreted for their isotopic light to heavy ratios. For those probe-modified peptides that showed ratios greater than two, we only interpreted those targets that were present across all three biological replicates, were statistically significant and showed good quality MS1 peak shapes across all biological replicates. Light versus heavy isotopic probe-modified peptide ratios are calculated by taking the mean of the ratios of each replicate paired light versus heavy precursor abundance for all peptide-spectral matches associated with a peptide. The paired abundances were also used to calculate a paired sample t-test P value to estimate constancy in paired abundances and significance in change between treatment and control. P values were corrected using the Benjamini–Hochberg method.

### GCLM Knockdown Studies

RNAi was performed by using siRNA purchased from Dharmacon specific to GCLM. A2780 cells were seeded at a density of 3x10^5 cells per mL in a 6-well format and allowed to adhere overnight. The next day, cells were transfected with 25 nM of small interfering RNA (siRNA)to GCLM (Dharmacon L-011670-01-0005) or a non-targeting negative control (ON-TARGETplus Non-targeting Control Pool, Dharmacon #D-001810-10-05) using 50 nM of DharmaFECT4 (Horizon Discovery, T-2005-01) as a transfection reagent. Transfection reagent was added to OPTIMEM (ThermoFisher, 31985070) media and allowed to incubate for 5 min at room temperature. Meanwhile, siRNA was added to an equal amount of OPTIMEM. Solutions of transfection reagent and siRNA in OPTIMEM were then combined and allowed to incubate for 30 minutes at room temperature. These combined solutions were diluted with complete DMEM to provide 200μl per well, and the media was exchanged. After 48 h, cells were supplemented with DMSO or compound and cultured for an additional 24 to 48 h before undergoing cell viability testing as described above. Cells were then harvested, and lysates were obtained to analyze protein abundance via Western blotting.

### Western Blots and Quantification

Proteins were resolved by SDS/PAGE and transferred to nitrocellulose membranes (BioRad, 1704159) using the Trans-Blot Turbo transfer system (Bio-Rad). Membranes were blocked with 5% BSA in Tris-buffered saline containing Tween 20 (TBST) solution for 1 hr at RT, washed in TBST, and probed with primary antibody diluted per recommended manufacturer procedures in 5% TBST and incubated while rotating overnight at 4°C. After 3 washes with TBST, the membranes were incubated in the dark with IR680- or IR800-conjugated secondary antibodies (LI-COR) at 1:10,000 dilution in 5% BSA in TBST at RT for 1 h. After 3 additional washes with TBST, blots were visualized using an Odyssey Li-Cor Clx fluorescent scanner. Antibodies used in this study were Rb to GCLM (Abcam, ab126704), Rb to GCLC (Abcam, ab190685), Ms to GAPDH (Proteintech Group Inc., 60004-1-Ig), IRDye 680RD Goat anti-Mouse (LI-COR 926-60870), and IRDye 800CW Goat anti-Rabbit (LICOR 926-32211). Band quantification was performed using ImageJ software.

### Ferrostatin Rescue Experiments

Cells were seeded in a 96 well plate at 10,000/well in a 200 μl volume and allowed to adhere overnight. Ferrostatin-1 (Cayman chemicals, 17729) was added to the wells (2μM final) at 2μl and followed with addition of 2μl of EN-25 (50μM final) or DMSO vehicle. Cells were incubated at 37°C for varying time points before proliferation was assayed by Hoechst staining.

## Author Contributions

LZ, DKN conceived of the project idea, designed experiments, performed experiments, analyzed and interpreted the data, and wrote the paper. MT, XT, QS, VT, SS, MK, YI, TI designed and performed experiments and analyzed and interpreted the data.

## Acknowledgement

We thank the members of the Nomura Research Group and Novartis Institutes for BioMedical Research for critical reading of the manuscript. This work was supported by Ono Pharmaceuticals for all authors. This work was also supported by the Nomura Research Group and the Mark Foundation for Cancer Research ASPIRE Award. We also thank Drs. Hasan Celik, Alicia Lund, and UC Berkeley’s NMR facility in the College of Chemistry (CoC-NMR) for spectroscopic assistance. Instruments in the College of Chemistry NMR facility are supported in part by NIH S10OD024998.

