## Supporting Information for "Covalent Targeting of Glutamate Cysteine Ligase to Inhibit Glutathione Synthesis"

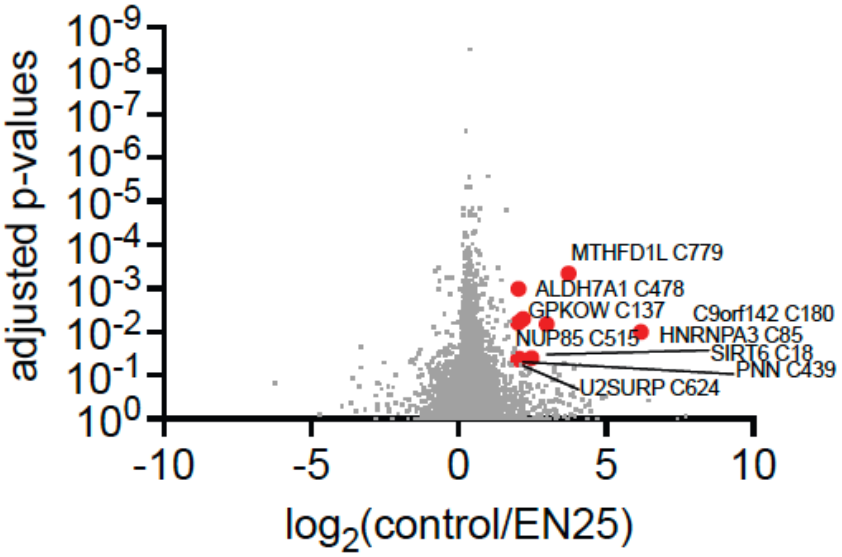


**Figure S1. isoTOP-ABPP analysis of EN25 in A2780 ovarian cancer cells.** A2780 cells were treated *in situ* with DMSO vehicle or EN25 (50 µM) for 90 min prior to labeling of the proteomes *ex situ* with IA-alkyne (100 µM) for 1 hr. Isotopically light (for DMSO-treated) or heavy (for compound-treated) TEV protease-cleavable biotin-azide tags were appended by CuAAC and probe-labeled proteomes were mixed in a 1:1 control/treated ratio, enriched with avidin, digested with trypsin, and eluted with TEV protease for isoTOP-ABPP analysis. Shown are control versus EN25 probe-modified peptide ratios and adjusted p-values for peptides identified in at least 2 out of the 4 biological replicates/group. Shown in red are the modified residues of targets that showed >2-fold control/EN25 ratio with adjusted p<0.05. Raw proteomic data is in **Table S2.**

**Supporting Table Legends**

**Table S1. Structures of covalent ligands screened against the GCL complex and percent inhibition values compared to DMSO controls using approaches described in Figure 1a.**

**Table S2. isoTOP-ABPP analysis of EN25 in A2780 ovarian cancer cells *in situ*.** A2780 cells were treated *in situ* with DMSO vehicle or EN25 (50 µM) for 90 min. DMSO and EN25 treated cell lysates were labeled with IA-alkyne (100 µM) for 1 h, after which isotopically light (control) or heavy (EN25-treated) biotin-azide bearing a TEV tag was appended by CuAAC. Proteomes were mixed in a 1:1 ratio, probe-labeled proteins were enriched with avidin and digested with trypsin, and probe-modified peptides were eluted by TEV protease and analyzed by LC-MS/MS. Data in table shows probe-modified peptides identified, the site of modification, the light to heavy probe-modified peptide ratio, and the protein identification from n=4 biological replicates per group.

### Synthetic Methods and Characterization

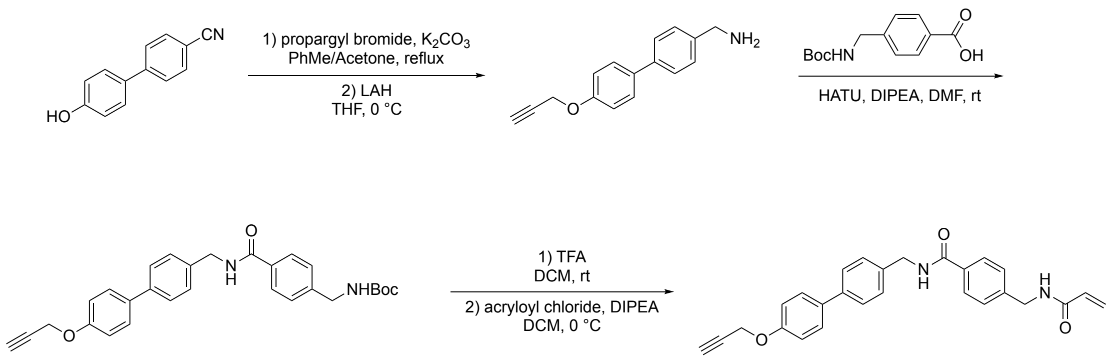


**Procedure for the preparation of EN25-alkyne probe:**

Treat the phenol compound with propargyl bromide under $K_2CO_3$ base condition in toluene and acetone solution. The reaction mixture was stirred for 6 hours under reflux. The product mixture was concentrated under vacuum and purified through column. LAH was added to the nitrile compound in THF at 0 $^oC$, stirred for 6 h, then purified through column.

Treat the amine compound with Boc protected benzoic acid with HATU and DIPEA in DMF, the reaction mixture was stirred overnight at room temperature. The product mixture was concentrated under vacuum and purified through column.

TFA was added to a vigorously stirring solution of the Boc protected alkyne compound in DCM. The reaction mixture was stirred for 2 hours at room temperature. The product mixture was concentrated under vacuum. Then acryloyl chloride and DIPEA was added to the compound in DCM at 0 $^oC$, stirred for overnight. The product mixture was concentrated under vacuum and purified through column.

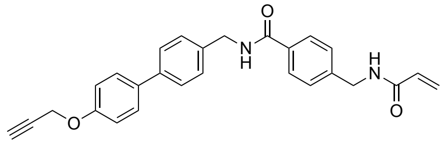

**4-(acrylamidomethyl)-*N*-((4'-(prop-2-yn-1-yloxy)-[1,1'-biphenyl]-4-yl)methyl)benzamide (EN25-alkyne probe)**

$^1$H NMR (500 MHz, DMSO) δ 9.02 (t, *J* = 6.0 Hz, 1H), 8.65 (d, *J* = 6.0 Hz, 1H), 7.87 (d, *J* = 7.9 Hz, 2H), 7.58 (t, *J* = 8.3 Hz, 4H), 7.36 (t, *J* = 7.8 Hz, 4H), 7.06 (d, *J* = 8.4 Hz, 2H), 6.29 (dd, *J* = 17.1, 10.2 Hz, 1H), 6.14 (dd, *J* = 17.1, 2.2 Hz, 1H), 5.64 (dd, *J* = 10.1, 2.2 Hz, 1H), 4.83 (s, 2H), 4.50 (d, *J* = 5.9 Hz, 2H), 4.41 (d, *J* = 5.9 Hz, 2H), 3.58 (s, 1H). $^{13}$C NMR (126 MHz, DMSO) δ 166.44, 165.15, 157.21, 143.15, 138.80, 138.71, 133.57, 133.43, 132.03, 128.28, 128.08, 127.83, 127.62, 126.65, 126.06, 115.76, 79.75, 78.78, 55.93, 42.80, 42.36. HRMS (ESI-TOF) Calcd for $C_{27}H_{24}N_2NaO_3^+$ $[M+Na]^+$ : 447.1679; found: 447.1683.

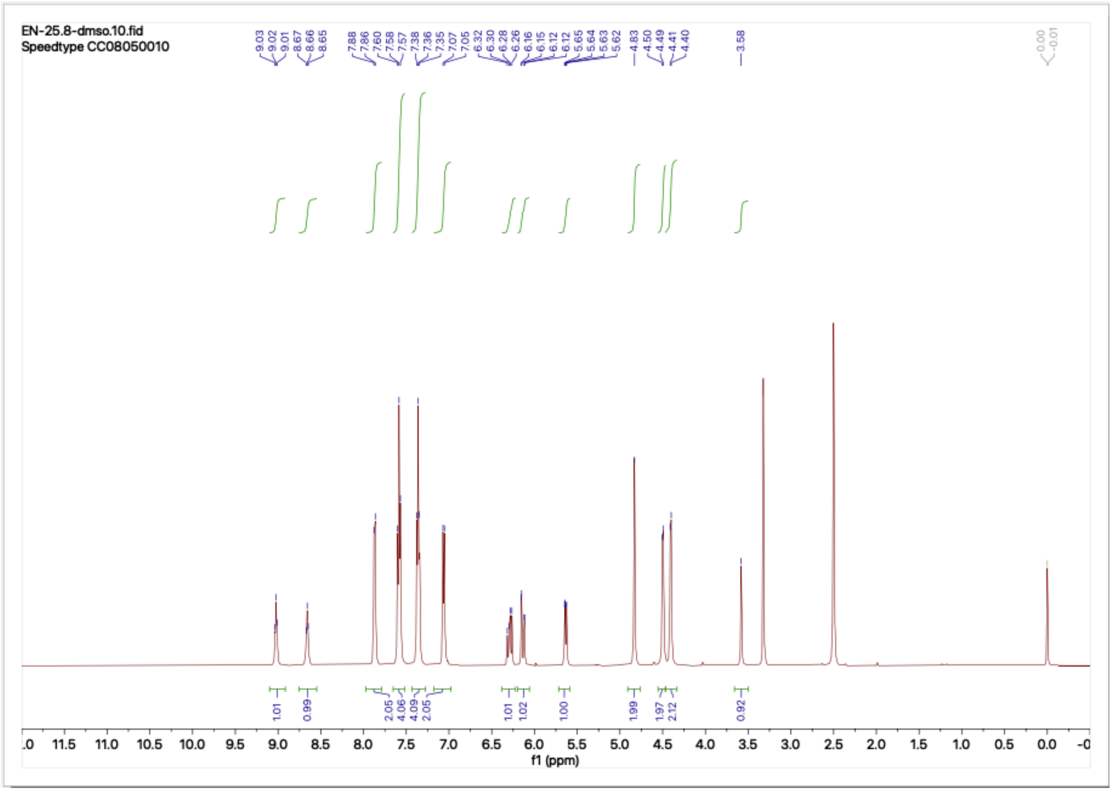

EN-25.8-dmso.10.fid
Speedtype CC08050010
9.03
9.02
9.01
8.67
8.66
8.65
7.88
7.86
7.60
7.58
7.57
7.38
7.36
7.35
7.07
7.05
6.32
6.30
6.28
6.26
6.16
6.15
6.12
6.12
5.65
5.64
5.63
5.62
4.83
4.50
4.49
4.41
4.40
3.58
0.00
-0.01
1.01
0.99
2.05
4.06
4.09
2.05
1.01
1.02
1.00
1.99
1.97
2.12
0.92
f1 (ppm)


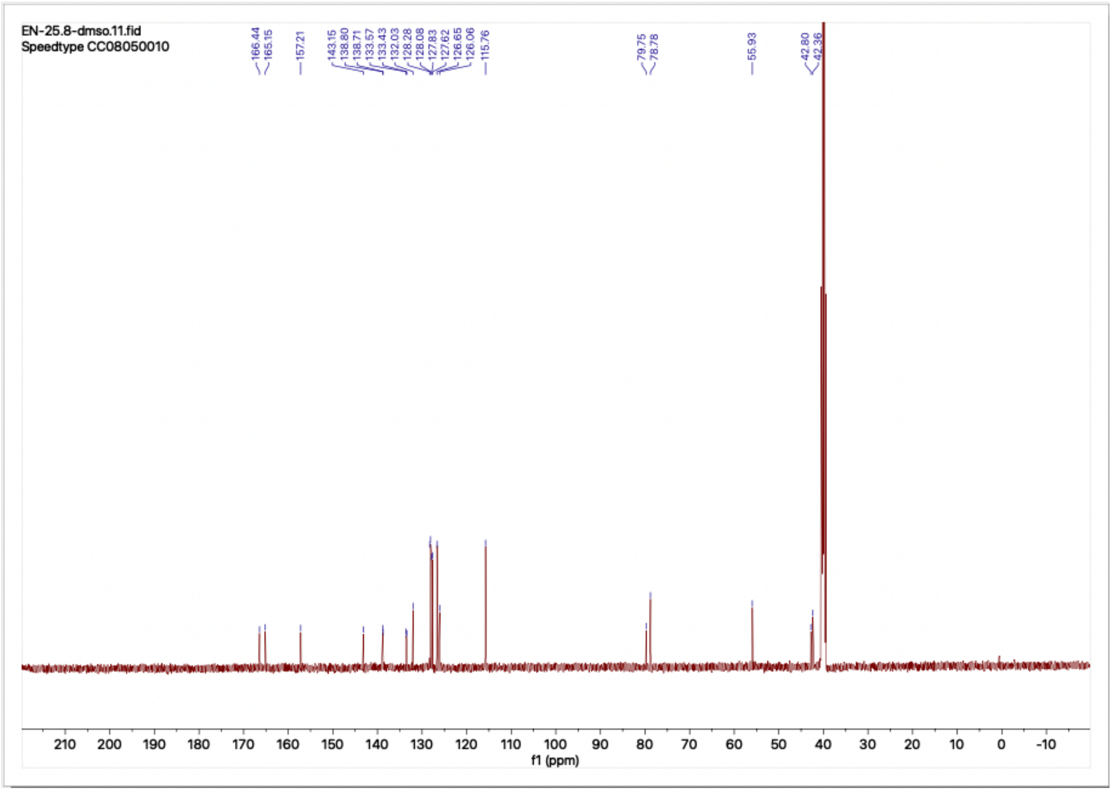

EN-25.8-dmso.11.fid
Speedtype CC08050010
166.44
165.15
157.21
143.15
138.80
138.71
133.57
133.43
132.03
128.28
128.08
127.83
127.62
126.65
126.06
115.76
79.75
78.78
55.93
42.80
42.36
f1 (ppm)
